# ICA treatment diabets induced bone loss via primary cilia/Gli2/Osteocalcin signaling pathway

**DOI:** 10.1101/2021.10.23.465584

**Authors:** Jie Liu, Xiangmei Wu, Xiaoyan Deng, Huifang zhu, Tingting Wang, Maorong Wang, Shengyong Yang, Jie Xu, Qian Chen, Mengxue Li, Xianjun Liu, Changdong Wang

## Abstract

Diabetes mellitus, as a metabolic system disorder disease, aggravates the disease burden of patients and affects the quality of human life. Diabetes-associated bone complications lead to decreased bone mechanical strength and osteoporosis. Evidences show that chronic hyperglycemia and metabolic intermediates, such as inflammatory factor, reactive oxygen species (ROS) and advanced glycation end products (AGEs), are regarded as dominant hazardous factors of primary cilia/Gli2 signal disorders. Case studies have demonstrated abnormal bone metabolism in diabetics, however, how diabetes damages primary cilia/Gli2 signal is largely unknown. Therefore, we studied the effects of diabetes on femoral primary cilia by establishing a Streptozocin (STZ)-induced diabetic (Sprague Dawley) SD rat model and diabetic bone loss cell model *in vitro*. Our results confirmed that diabetes impaired femur primary cilia, osteoblast differentiation and mineralization by inhibiting primary cilia/Gli2 signaling pathway, additionally, Icariin(ICA) treatment could rescue the impairment of osteoblast differentiation caused by high glucose medium *in vitro*. ICA activated primary cilia/Gli2/osteocalcin signaling pathway of osteoblasts by protecting primary cilia from glucotoxicity imposed by diabetes, intact primary cilia could be as anchoring sites, in which Gli2 was processed and modified, and matured Gli2 entered the nucleus to initiate downstream osteocalcin gene transcription. Additionally, ICA inhibited ROS production of mitochondria, thus balanced mitochondrial energy metabolism and oxidative phosphorylation. All results suggest that ICA can protect the primary cilia and mitochondria of osteoblast by reducing intracellular ROS, thereby recover primary cilia/Gli2 signaling pathway to facilitate osteoblast differentiation and mineralization, suggesting that ICA has potential as a novel type of drug treating bone loss induced by diabetes.

## Introduction

Diabetes (diabetes mellitus, DM) is a global disease which severely threatens human health after cardiovascular diseases and tumors. According to statistics, the number of diabetic patients has reached about 8%-9% (450 million) of the world’s total population. The prevalence rate has been increasing year by year and reached epidemic level, and the age of patients is getting younger. There are about 110 million diabetic patients in China, ranking first in the world^[1, 2]^. Compared with healthy people of the same age, diabetic patients have two to ten times higher risk of death due to cardiovascular diseases such as coronary heart disease and stroke, and the risk of suffering from retinopathy and bone microvascular complications is also increased^[3]^.

More than 9 million osteoporotic fractures occur worldwide each year, among which diabetes is an important risk factor for the increase in the global incidence of osteoporosis^[4]^. The continuous remodeling of the bone cycle depends on the balance between osteogenesis and bone resorption. Diabetic patients appear decreased bone healing and regeneration capabilities and increased fracture risk^[5]^. The pathophysiological mechanism that induces bone fragility in diabetic patients is fairly complex, including the accumulation of inflammatory factors, ROS and AGEs, which destroy the collagen structure, increase the bone marrow fat content, and change the function of bone cells^[6]^. Futhermore, changing the ratio of OPG/RANK/RANKL breaks the balance mechanism of bone remodeling^[7]^. Recent studies have found that iron overload and ROS accumulation in type 2 diabetes lead to mitochondrial dysfunction and ferroptosis in osteoblasts^[8]^.

A group of reactive molecules and free radicals, including superoxide, hydrogen peroxide and hydroxy radical et., were collectively termed as ROS^[9]^, it is an important regulator in the metabolic reaction process of living organism, not only can regulate the redox balance, but also a signal molecule ^[10]^. ROS keep ambivalent because it is both essential and detrimental to life, the tipping point depends on concentration. Currently, cumulative evidences have confirmed that long-term elevated blood glucose will lead to excessive ROS production in mitochondria, resulting in damage to downstream cells and organs^[11]^, which suggests the possibility of mitochondrial-targeted ROS’s therapy of diabetic complications to inhibit downstream deleterious metabolic pathways.

The bone mineral density (BMD) of type 1 diabetes mellitus (T1DM) patientsis significantly decreased, yet there’s a controversy that BMD of type 2 diabetes mellitus (T2DM) patients is not significantly reduced or even increased, but its bone transformation efficiency is very low^[12]^. However, both T1DM and T2DM, there will be a downtrend in bone mechanical strength but an increased risk of fracture when the patient is not adequately treated^[13]^. Current research proves that osteoporosis can be prevented and treated, therapeutic agents have evolved from initial estrogen and calcitonin to current bisphosphonates^[14]^, targeted nuclear factor kappa B receptor activator ligand (RANKL) (Denosumab)^[15]^, parathyroid hormone analogue (Teriparatide)^[16]^, and targeted sclerostin monoclonal antibody (Romosozumab) ^[17]^. Although remarkable progress has been achieved in the development of therapeutic drugs, we still need to pay attention to the side effects and long-term effects of these drugs. Adverse reaction report indicates that Teriparatide has the risk of osteosarcoma^[18]^, and bisphosphonates cause atypical femur fractures and osteonecrosis, and the efficacy after 5 years of treatment has not been determined^[19, 20]^. Therefore, in order to alleviate patients’ worries about drugs so that they can get seasonable treatment, we need to find therapeutic drugs from other fields to inhibit the bone resorption of osteoclasts and enhance the osteogenesis of osteoblasts. Traditional Chinese medicine *Epimedium* has long been used to treat fractures and prevent osteoporosis^[21]^, Icariin (ICA), a prenylatedflavonol glycoside extracted from *Epimedium*. ICA can relieve the inhibition of PDE5 on aromatase P450 and promote the catalytic synthesis of estrogen from C19 androgen^[22, 23]^. Animal experiments have shown that ICA can act as an estrogen analog to protect ovariectomized rats by increasing osteogenesis and angiogenesis^[24]^, the underlying mechanisms is that ICA can activate ERα and Akt by inducing IGF-1 production to promote bone formation ^[25]^. In addition, cell experiments have shown that ICA can increase the osteoblast differentiation and mineralization of bone marrow mesenchyml stem cells (BMSCs), and inhibit the formation of osteoclasts and bone resorption^[26–28]^.

Cilia are mainly divided into motile cilia and primary cilia, the latter originate from the mother centrioles of the G0/G1 stage of the cell. In evolution, as a strictly conserved information transmission hub organelle, primary cilia are similar to an “antenna” protruding from the plasma membrane, the presence of primary cilia has been observed in osteocytes, osteoblasts, and chondrocytes^[29]^. And as a mechanical sensor of bone, primary cilia play an important role in the proliferation, differentiation and morphological maintenance of bone cells^[30]^. However, the primary cilia of osteocyte are sensitive organs for mechanical stimulation, abnormal cilia morphology has been observed in bone diseases such as short rid-polydactyly syndrome (SRPs) and Jeune asphyxiating thoracic dystrophy (JATD). *Oliver Kluth* et al. found that the number of primary cilia of pancreatic islet cells in diabetic rat models was significantly reduced^[31]^, which may be related to the high levels of inflammatory factors and ROS ^[32]^. Prolonged period of high levels of TGF-β in diabetic patients can induce the expression of HDAC6, which deacetylates acetylated α-tubulin leading to cilia shortening and even degradation, thereby preventing osteoblasts from maturation^[33]^, suggesting that primary cilia can respond to elevated blood glucose by deassembly. Primary cilia integrate islet cell crosslinking and glucose homeostasis regulation, specific deletion of islet β cell cilia not only impairs insulin secretion, but also affects glucagon and somatostatin secretion of proximal α and δ cells, leading to diabetes^[34]^. Studies by *Shi* et al. confirm that ICA can enhance cAMP/PKA/CREB signals in the primary cilia to promote the maturation and mineralization of osteoblasts^[35]^.

Here, We explored case data and confirmed that large accumulations of lipid in the serum of diabetic patients were threats to diabetes induced bone loss, and that inorganic mineral ions Ca^2+^ related to bone metabolism, ALP and glycated hemoglobin A1c (HbA1c) are closely related. In the SD rat diabetes model, abnormal lipid metabolism and glycogen metabolism occurred, the microstructure of the femur was changed, and the bone biomechanical properties were reduced. Further studies confirmed that the number of femoral primary cilia was decreased significantly, which was linked to the excess of ROS caused by diabetes. *In vitro* cell model, ICA could act as a ROS inhibitor to rescue osteoblasts differentiation and mineralization inhibition rendered by high glucose by activating primary cilia/Gli2 pathway. We will continue to study the mechanism of ICA as a drug on diabetic bone complications.

## Results 1. The serum lipid, ALP and Ca^2+^ reflect bone metabolism disorders

To investigate the effect of diabetes on bone development, the date of 399 patients of T2DM (male:261, female:138) were collected to value the association between gender, age, serum mineral ions, lipid, ALP and osteoporosis. Case inclusion criteria were : male and female between the ages of 20 and 75 who have been diagnosed with T2DM for more than 1 year. The pathophysiological mechanisms that induce bone weakness in diabetic patients are very complex, including hyperglycemia, oxidative stress and accumulation of AGEs, which can destroy the properties of collagen, increase the content of bone marrow fat, and possibly change the function of bone cells^[6]^. The results of binary Logistic regression showed that high HDL, aging, female and low serum Ca^2+^ were risk factors of osteoporosis (Figure 1A). Chronic hyperglycemia leaded to a toxic internal environment when diabetes couldn’t be effectively treated, Pearson correlation analysis results revealed that the correlations between HbA1c and serum apolipoprotein B (ApoB), alkaline phosphatase (ALP) are positive (r=0.256, r=0.427, respectively), the correlation between HbA1cand serum Ca^2+^ is negative (r=-0.272) (Fig. 1B). Subsequently, with HbAc1 as the outcome variable, ApoB, Ca^2+^, ALP as the independent variables, a multiple linear regression was performed, the results proved ApoB, ALP and Ca^2+^ had impacts on HbA1c (R^2^=0.286, P < 0.001)(Fig. 1C). Clinical data suggests that we should focus more on serum lipid levels and mineral ion levels, and keeping HbA1c at a healthy levels is important for bone metabolism.

**Figure 1:**
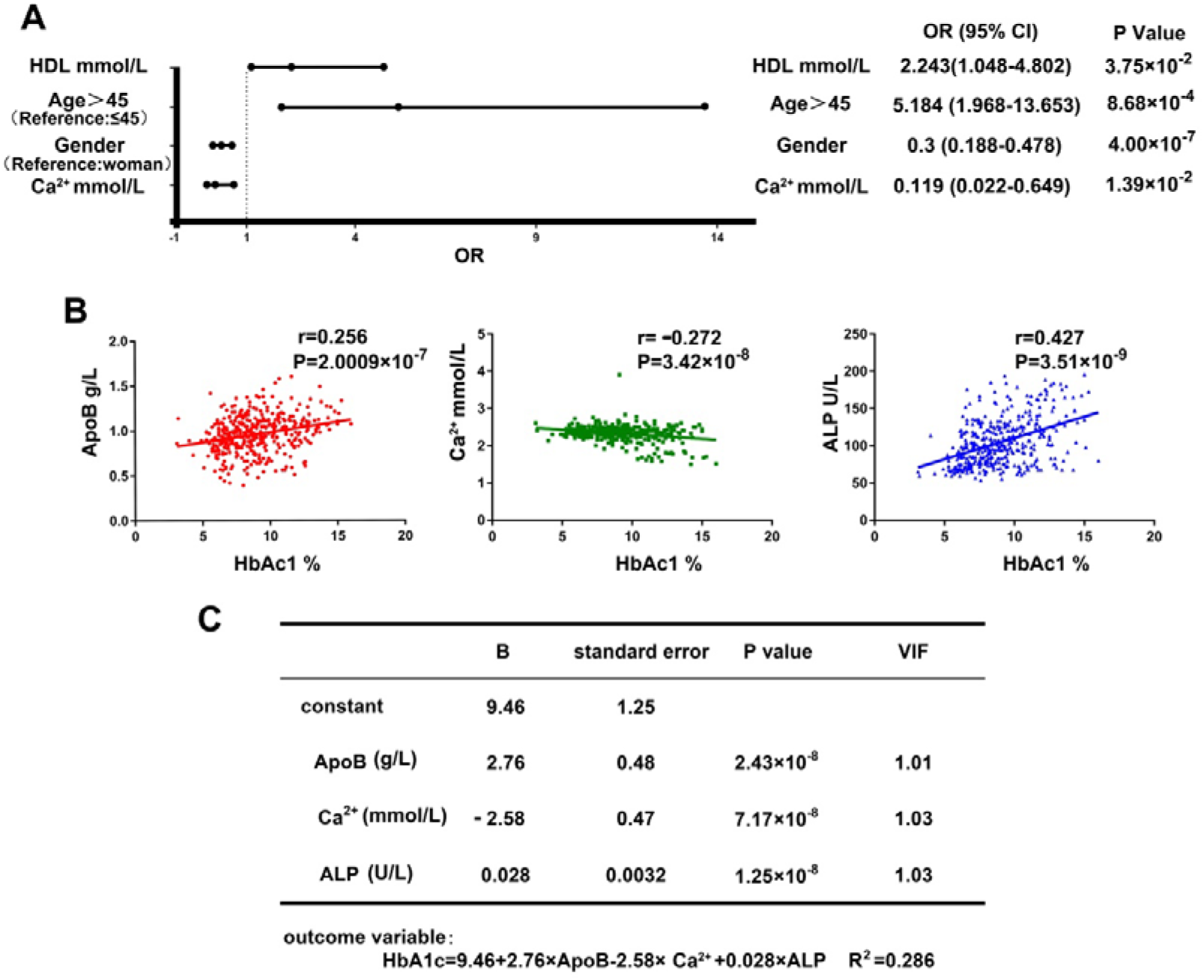
Case data from diabetic patients. (A) Binary Logistic regression analysed risk factors of osteoporosis. (B) Pearson correlation analysis between HbA1c and serum ApoB, Ca^2+^, ALP, (C) the model of multiple linear regression of HbA1c, serum ApoB, Ca^2+^ and ALP.

## Results 2. Intraperitoneal injection of high dose STZ damage pancreatic islet β-cells causing metabolic disorders and diabetic phenotype in SD rats

Diabetes leads to decreased bone mechanical strength and increased fracture risk^[13]^. To study the mechanism of diabetic bone disease, we first established a T1DM SD rats model by intraperitoneally injecting at a single large dose STZ ^[36]^. Compared with vehicle control group, we found that high-dose STZ injection leading to a diabetic phenotype, such as losing weight (Figure 2A) and prolonged period of hyperglycemia (Figure 2B) in SD rats. The reason was that STZ had toxic effects on pancreatic islet β-cells specially, and the number and function of pancreatic islet cells were closely related to energy metabolism, a single high-dose STZ modified DNA by alkylation caused almost all pancreatic β-cell necrosis^[37, 38]^. We performed H&E staining on pancreatic sections and the results showed that the structure and morphology of the pancreas of SD rats in the vehicle control group were intact, but the cells appeared deterioration, lysed nuclei and enlarged intercellular spaces in diabetes group (Figure 2C). The organism dysfunction induced by diabetes caused disability to respond or adapt to changes of metabolic demand, whereby cannot utilize glucose and fatty acids flexibly^[39]^. We performed transmission electron microscopy to observe microstructure of tissue cells of pancreas and spleen from vehicle control and the STZ-induced diabetic SD rats, respectively. The results showed that a large number of glycogen particles could be seen in the spleen tissue of STZ-induced diabetic SD rats, while there was no glycogen particles in vehicle control group (Figure 2D, blue box). Meanwhile, compared with vehicle control group, lipid droplets accumulation (Figure 2E, red box) and swelling mitochondria (Figure 2E, black box) were observed in pancreatic tissue of diabetic SD rats. These results confirm that high-dose injection of STZ destroys the structure and function of the pancreatic islet β-cell, which contributes to developed hallmark features of diabetes, such as metabolism disorder of glucose and lipid, as well as low body weight. Diabetic complications involve various organs in the organism, such as the kidney, brain and retina^[40, 41]^, and insulin deficiency causes energy metabolism disorders.

**Figure 2.**
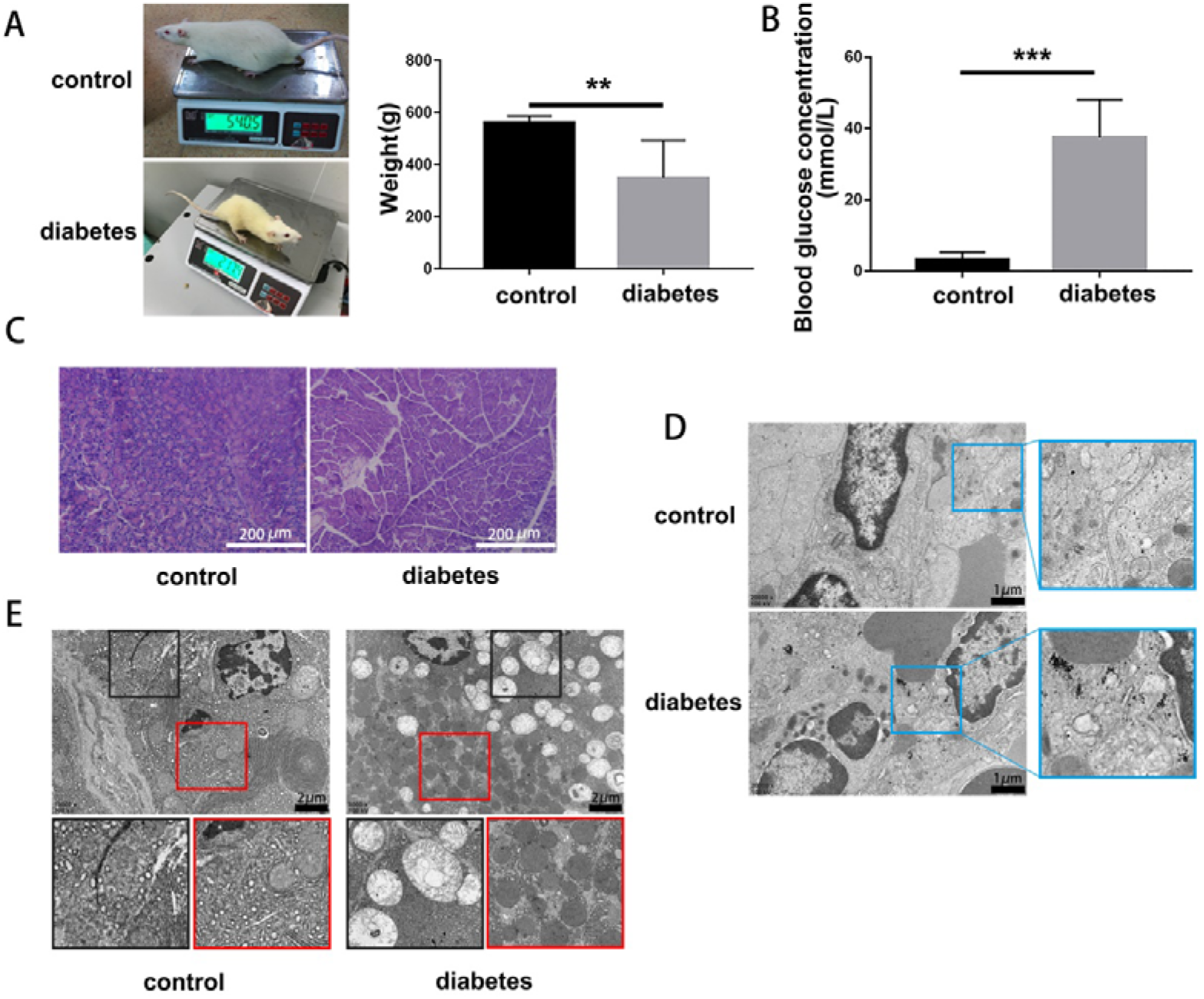
STZ-induced SD rats occur symptoms related with diabetes. To study the effect of hyperglycemia on bone metabolism, we established STZ-induced diabetic SD rats model and fed them for 90 days. (A) Weigh and photograph SD rats in the STZ group and the vehicle control group. (B) Serum was collected from the tail vein to detect glucose concentration of the two groups of SD rats. (C) The pancreatic tissue sections were subjected to H&E Staining (scale bar: 200μm). (D)Transmission electron microscope observed of spleen tissues (scale bar: 1μm, blue box: glycogen particles) and (E) pancreatic tissues (scale bar: 2μm, red box: lipid droplets, black box: mitochondria) and Data were expressed as mean±SEM and were processed by SPSS.22 Student’s t test. *P<0.05, **P<0.01 and ***P<0.001, compared with vehicle control SD rats.

## Results 3. Bone loss induced by diabetes results in severe impairment of bone mechanical properties in SD rats

Superior bone mechanical properties are the basis for bones to support movement and protect important organs^[42]^. To explore the effects of diabetes on the bone microarchitecture, we utilized Micro-CT, an apparatus with extremely high resolution, to observe the internal structure of bone in the premise of maintaining the integrity of the bone structure^[43, 44]^, following assessed both cortical and trabecular bone morphology and microstructure of model SD rats after fed for 90 days. The related parameters of total tissue volume (TV), bone volume (BV), bone volume fraction (BV/TV) reflect bone mass and bone metabolism. Compared with the control group, Micro-CT results showed that diabetic SD rats TV, BV and BV/TVwere significantly reduced (Figure 3A, 3B). Bone trabecular number (Tb.N), trabecular thickness (Tb.Th) and trabecular separation (Tb.Sp) are main parameters to evaluate the spatial structure of bone trabecula. Micro-CT results proved that Tb.N and Tb.Th reduced notably, while Tb.Sp increased significantly (Figure 3B). Bone is usually divided into compact cortical bone and lattice-like arrangement of cancellous bone. Cortical bone is mainly pronounced within the diaphysis, while cancellous bone is mainly distributed in the metaphysis and epiphysis^[45, 46]^. The biomechanical detection of femur found that the mechanical strength of cortical bone (figure 3C, 3D) and cancellous bone (figure 3E, 3F) were significantly weakened, confirming that diabetes caused damage to the mechanical properties of bone. Bone fulfills functions in maintaining blood calcium levels, provides support and protection for soft tissues^[47]^, and it’s also an endocrine organ^[48]^.The osteocalcin(OCN) secreted by bone is not just a marker of osteoblast differentiation, and can regulate glucose metabolism^[49]^. We used an ELISA kit to detect the OCN concentration in the femur protein and serum of model SD rats and found that the OCN concentration was significantly reduced in diabetic SD rats (Figure 3G). Alkaline phosphatase (ALP) catalyzes hydrolysis of phosphomonoesters and promotes the formation of hydroxyapatite and the mineralization of the bone matrix^[50, 51]^, serum ALP mainly derives from bone and liver^[52]^. We used an ELISA kit to detect the ALP activity and found that, compared with control group SD rats, the ALP activity of femoral protein from diabetic model SD rats was significantly reduced, while serum ALP activity was significantly increased (Figure 3H), abnormal OCN concentration and ALP activity were signs of bone diseases. H&E staining performed on the distal tibia tissues showed that diabetes caused distinct osteopenia (Figure 3I). To more rigorously examine bone growth status, femur protein was used to detect the expression of osteoblast differentiation maker type I collagen (COL □), runt-related transcription factor 2 (Runx2), osteopontin (OPN), ALP, OCN by Western blot and results showed that the expressions of COL □, Runx2, OPN, ALP and OCN reduced significantly (Figure J, K). These results confirm that the osteoblast differentiation inhibited by diabetes is one of the pivotal causes of bone loss, which eventually damages to the microstructure and mechanical properties of cortical bone and trabecular bone.

**Figure 3.**
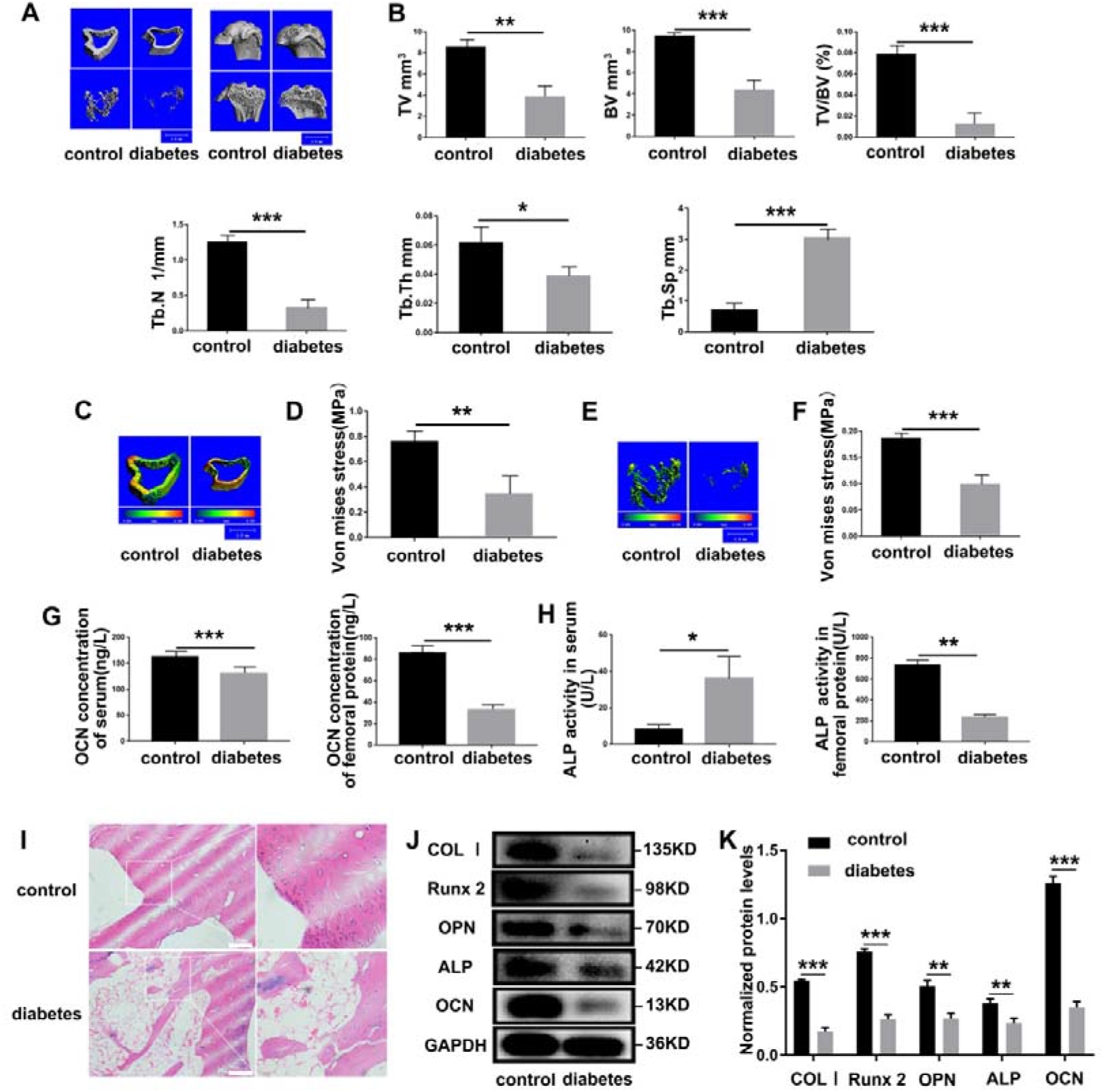
Diabetes cause severe bone defects. (A) Micro-CT showed the results of transverse section and longitudinal section of the distant femur. (B) Quantitative analysis Micro-CT-derived bone microstructural parameters of total tissue volume (TV), bone volume (BV), bone volume fraction (BV/TV), trabecular bone number (Tb.N), trabecular bone thickness (Tb.Th) and trabecular bone separation (Tb.Sp). (C) Micro-CT showed a comparison of the stress levels of the cortical bone. (D) Quantitative analysis Micro-CT-derived bone stress levels of (C). (E) Micro-CT showed a comparison of the stress levels of the cancellous bone. (F) Quantitative analysis Micro-CT-derived bone stress levels of (E). (G) ELISA kit assayed serum and femur OCN concentration in rats. (H) ELISA kit assayed serum and femur ALP activity in rats. (I) H&E staining performed on the distal tibia tissues. (J) Western blot analysis of the protein expressions of COLⅠ, Runx2, OPN, ALP and OCN in femur. (K) Quantitative analysis the expression levels of (J). Data were expressed as mean±SEM and were processed by SPSS.22 Student’s t test. *P<0.05, **P<0.01 and ***P<0.001, compared with vehicle control SD rats.

## Result 4. ICA rescue diabetic bone loss *in vitro* cell model

To study whether high concentration of glucose affects the differentiation and mineralization of osteoblast, we refered to the method of Hernández et al. ^[53]^ and utilized three different osteoblast cell lines derived from rat bone marrow mesenchymal stem cells (BMSC), osteoblasts derived from newborn rat calvarian (primary OB) and mouse embryonic osteoblast precursor cells (MC3T3-E1) to constructed cell model of diabetes *in vitro*, by the means of adding D-(+)-glucose to medium to simulate a high glucose environment. In addition, we explore Icariin (ICA), which was extracted from the traditional Chinese medicine *Epimedium* and had therapeutic effect on fractures and osteoporosis^[21]^, whether could rescue osteoblast differentiation and mineralization in high glucose environment. We have determined optimal concentration of ICA is 40μmol/L through preliminary experiments, which had best effects on osteogenesis. The glucose concentration in control group (NC group) was 5.6 mM and in high glucose group (HG group) and ICA treatment group (HG+ICA) was 25 mM, respectively. The mineralization process of extracellular matrix was a unique characteristic of vertebrate bone system development. Carbonated apatite particles, together with collagen type I, water and non-collagen proteins collaboratively constituted the mineral phase of bone^[54]^, Alizarin red could specifically stain Ca^2+^-rich substances, so we induced three osteoblast cell lines by osteoblast differentiation medium with different glucose concentrations for 21 days, and alizarin red staining detected mineralization. Alizarin red staining results showed that the formation of mineralized nodules in the HG group was significantly less than that in the NC group, addition ICA significantly increased mineralization in HG+ICA group (Figure 4A, B, C). Meanwhile, we induced three osteoblast cell lines by osteoblast differentiation medium with different glucose concentrations for 3 days to extract total cell protein, ELISA kit assayed osteoblast differentiation marker ALP activity. The results showed that the ALP activity in the HG group was significantly lower than that in the NC group, while in HG+ICA group, addition ICA significantly increased ALP activity (Figure 4D, E, F). Then, MC3T3-E1 total cell protein was detected by western blot to analyse COL □ and OPN expression, compared with NC group, the results showed that COL □ and OPN expression was significantly reduced, surprisingly, adding ICA to 25mM high-glucose medium (HG+ICA group) markedly increased expression of these proteins (Figure 4G, H). Therefore, we successfully constructed cell models of diabetic bone loss *in vitro*, and found that ICA could be used as a drug to treat diabetic bone loss induced by high glucose. This *in vitro* model provides fundamental insights into ICA and facilitates our subsequent exploration about the medicinal mechanism of ICA treatment for bone loss.

**Figure 4.**
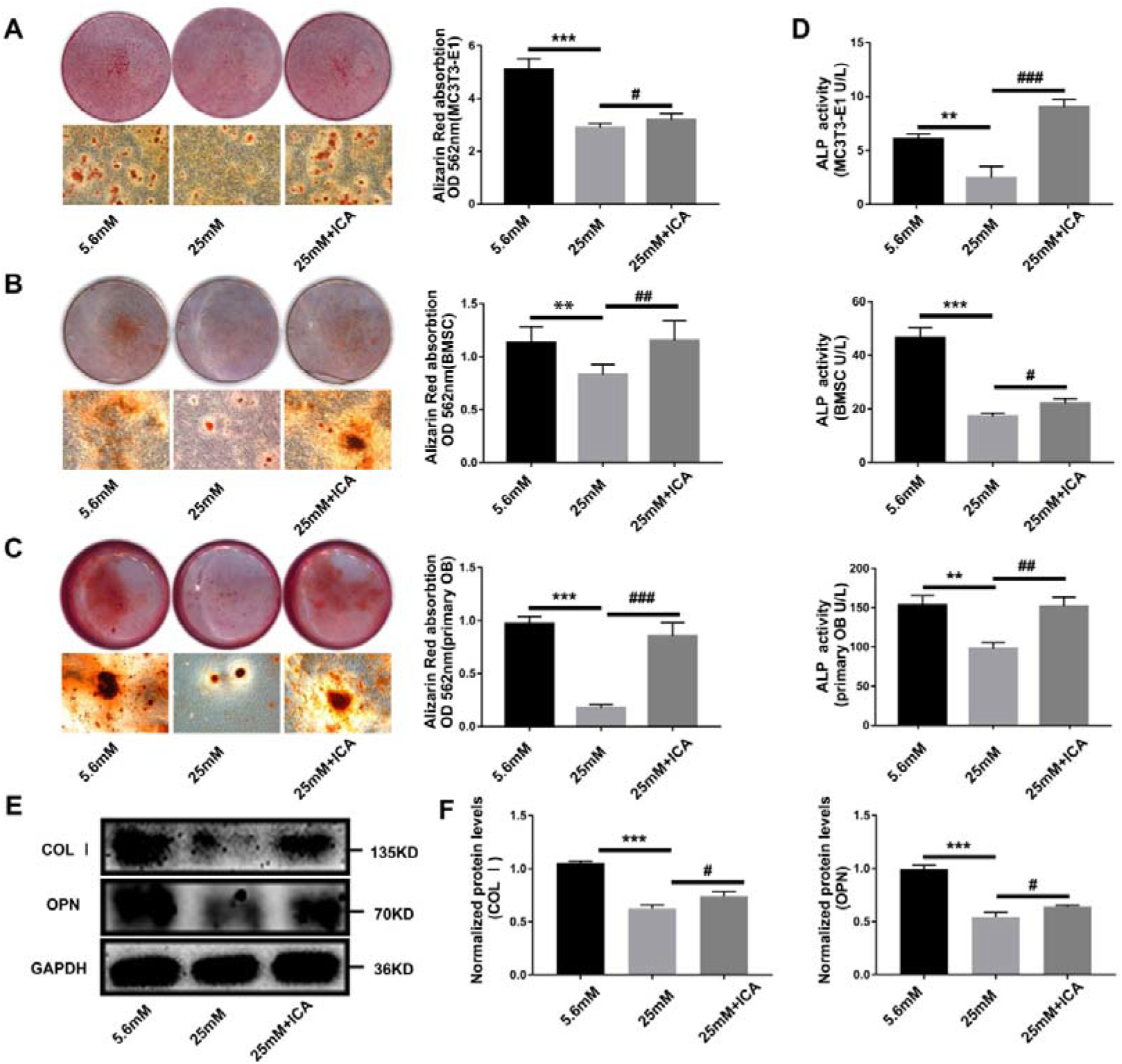
ICA plays a therapeutic role in a diabetic bone loss cell model. Three different osteoblast cell lines were induced in NC group, HG group and HG+ICA group for 21 days, and the extracellular matrix mineralized nodules were stained with Alizarin Red. The results of Alizarin red staining results and quantitative analysis of MC3T3-E1 (A), BMSC (B) and primary OB (C) cell. Three different osteoblast cell lines were induced in NC group, HG group and HG+ICA group for 3 days, extracted total cell protein to analyze by ELISA kit and western blot. (D) ALP activity assay of total cell protein of MC3T3-E1, BMSC and primary OB cell. (E) Western blot analyzed the expression of COL Ⅰ and OPN. (F) Quantitative analysis the level of protein expression in (D). Data were expressed as mean±SEM and were processed by SPSS.22 Student’s t test and one-way ANOVA. *P<0.05, **P<0.01 and ***P<0.001, compared with NC group. ^#^P<0.05, ^##^P<0.01 and ^###^P<0.001, compared with HG group.

## Results 5. Diabetes impairs primary cilia of splenic tissue and femur tissue, ICA can promote the growth of the osteoblast primary cilia and protect them from high glucose damage

Primary cilium is a kind of signal hub organelle, which can play a vital role as bone mechanical sensor. There are numerous specific receptors and ion channel proteins in ciliary membrane. It can sense osmotic pressure, hydrostatic pressure, Ca^2+^ flux and other physical stimuli and regulate paracrine through signaling pathways such as Hedgehog, Wnt, cAMP-PKA /CREB^[55]^, thereby regulating osteoblast proliferation, differentiation and matrix deposition^[56]^. Oliver Kluth et al. found that the number of primary cilia in pancreatic islets of diabetic rat models was significantly reduced, suggesting that primary cilia could disassemble in response to hyperglycemia^[57]^.The pancreatic tissue of SD rats were subjected to immunofluorescence and the results confirmed that, compared with the vehicle control group, almost all the primary cilia missed in diabetic SD rats (Fig. 5A), and the average fluorescence intensity of ciliary structural proteins acetylated α-tubulin (Ac-α-tubulin) and γ-tubulin were significantly reduced (Fig. 5B). Primary cilia integrated multiple effects on islet cells cross talk and glucose homeostasis, specifically removing primary cilia of pancreatic β cells not only impairs insulin secretion, but also affects glucagon and somatostatin secretion of α- and δ-cells, leading to diabetes mellitus^[34]^. So we hypothesized that diabetes would impair primary cilia of bone tissue. Through collecting the femurs of SD rats from vehicle control group and diabetes group for immunofluorescence. The results showed that, compared with vehicle control group, the femoral tissue of diabetic SD rats showed few primary cilia (Figure 5C), quantitative analysis results further confirmed the average fluorescence intensity of Ac-α-tubulin and γ-tubulin decreased significantly (Figure 5D). Western blot detected femur protein also found that the expressions of Ac-α-tubulin and γ-tubulin protein were significantly decreased (Figure 5E, F). These results implied diabetes damages the primary cilia on the pancreas, as well as the primary cilia on bone tissue and other tissues. Studies by Shi et al. confirmed that ICA could enhance cAMP/PKA/CREB signals in the primary cilia to promote the maturation and mineralization of osteoblasts^[35]^. Our study was designed to explore whether ICA could promote osteogenesis by protecting the primary cilia from high glucose damage. We cultured MC3T3-E1 cell in high glucose environment and observed the changes of primary cilia. By immunofluorescence, we found ICA promoted primary cilia growth and rescued primary cilia inhibited by high glucose (Figure 5G). Contrasted with NC group, Western blot results showed that the expression of primary cilia structural proteins Ac-α-tubulin and γ-tubulin were hindered significantly by high glucose, however, ICA could restore the expression of primary cilia structural proteins effectively (Figure 5H). Based on these results, we speculate that ICA has therapeutic effects on bone loss caused by diabetes, likely by protecting osteoblast primary cilia, thereby the primary cilia can normally fulfill function of information hub to regulate the process of bone formation.

**Fig 5.**
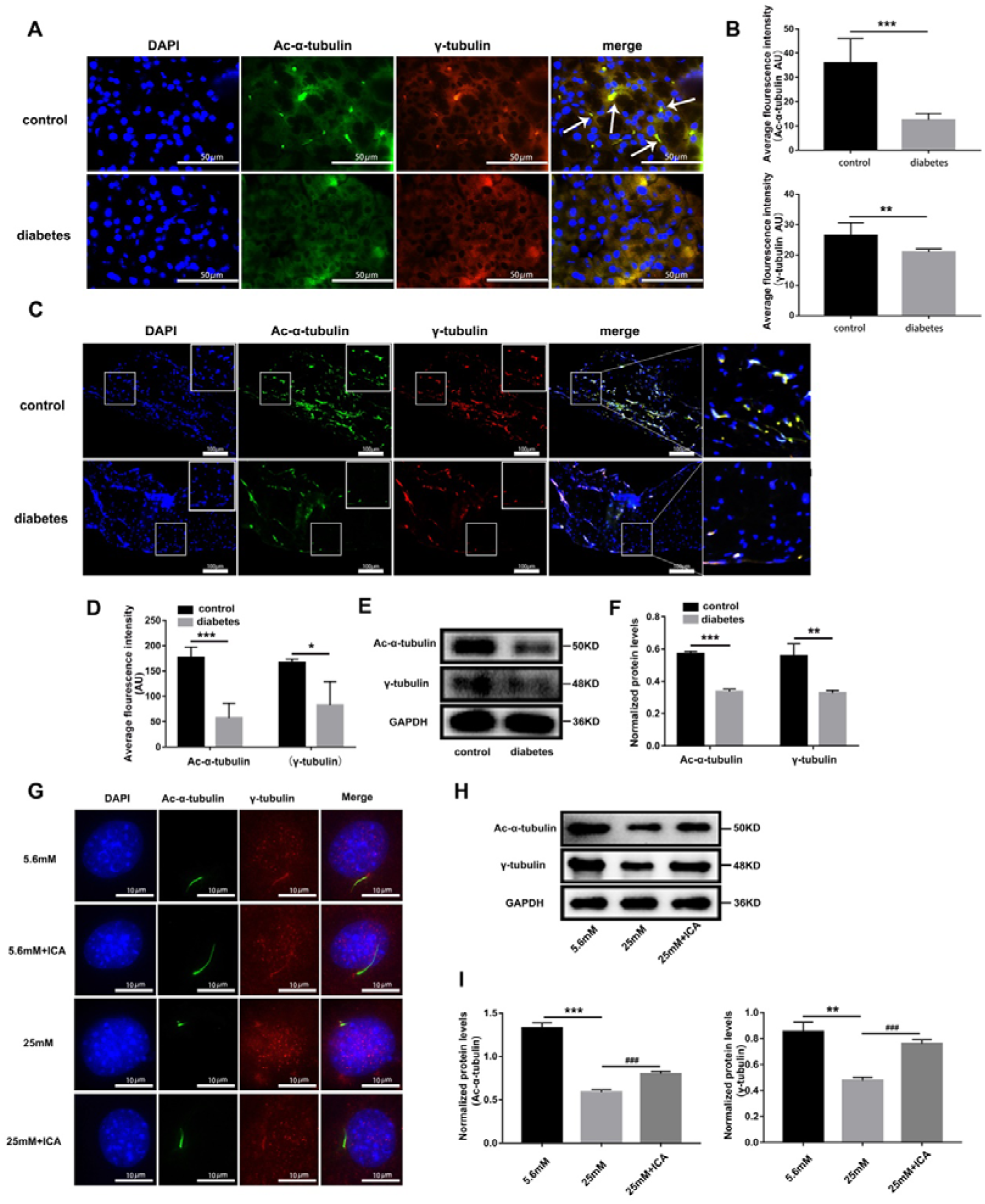
Diabetes impaire primary cilia in bone tissue and pancreas tissue, ICA could protect the primary cilia from high glucose damage. (A) Immunofluorescence of pancreas tissue, scale bar: 50μm, (B) quantitative analysis average fluorescence intensity of Ac-α-tubulin and γ-tubulin in (A). (C) Immunofluorescence of femur tissue, scale bar: 100μm, (D) quantitative analysis average fluorescence intensity of Ac-α-tubulin and γ-tubulin in (C). (E) Western blot detected the expression of Ac-α-tubulin and γ-tubulin in the femoral protein, (F) Quantitative analysis the level of protein expression in (E). ICA was added to the diabetic bone loss cell model and induced for 3 days, (G) immunofluorescence observed primary cilia. (H) Extracted cell total protein, western blot detects protein expression of Ac-α-tubulin and γ-tubulin. (I) Quantitative analysis the level of protein expression in (H). Data were expressed as mean±SEM and were processed by SPSS.22 Student’s t test and one-way ANOVA. *P<0.05, **P<0.01 and ***P<0.001, compared with NC group. ^#^P<0.05,^##^P<0.01 and ^###^P<0.001, compared with HG group.

## Results 6: ICA can protect anchor site of Gli2, activate Hedgehog signal and promote osteoblast differentiation

Hedgehog signaling is essential for the development, homeostasis, and regeneration of almost all organs in mammals, and downstream functions of PTCH1 and SMO depend entirely on cilia^[58]^. Three ligand molecules, SHH, IHH and DHH, respectively constitute different Hedgehog signaling networks to drive cell fate decisions and regulate the expression of target genes. SHH plays a particularly significant role in the development of nervous system and limbs, IHH plays a key role in the regulation of skeletal formation, furthermore, DHH plays a part in gonad development^[59]^.We detected the expression of Hedgehog signaling ligand molecules in the diabetic bone loss cell model, and the results showed that compared with the NC group, the expression of Gli2, SHH in the HG group was significantly reduced, and the expression of Ptch1, Sufu was significantly increased (Figure 6A, 6B). However, after addition ICA to high glucose medium, the expression of Gli2, SHH and IHH was significantly increased in recovery, meanwhile, the expression of Ptch1, Sufu were reduced remarkably (Figure 6A, 6B). Above results proved high glucose hampered Hedgehog signal by regulating signal molecular production. Gli protein can be proteolytically processed into three nuclear transcription factors——Gli1, Gli2 and Gli3, which responded to upstream Hedgehog ligand molecules. The current studies have shown that when absence of Hedgehog ligand, Gli protein remains a full-length unprocessed state and locates on the top of the primary cilia^[60]^. Immunofluorescence was used to detect the localization of Gli2 on the primary cilia under different glucose concentrations. In the NC group, Gli2 was co-localized with the ciliary axon protein Ac-a-tubulin, while in the HG group, the Gli2 disappeared and primary cilia shortened (Figure 6C, 6D), this suggests that the deletion of primary cilia may lead to the loss of anchor sites and degradation of Gli2. Consistent with previous results, addition ICA could protect the primary cilia from the damage of high glucose environment, and also rescue the expression of primary cilia and Gli2 (Figure 6C, 6D). To further determine whether ICA can restore Gli2 expression by protecting cilia, we used Cyclopamine (CYC), specific inhibitor of Hedgehog pathway, coupled with ICA to treat cells (HG +ICA+ CYC group). Compared with the HG+ICA group, the addition of CYC resulted in suppression of Gli2 expression even though the primary cilia expressed normally (Figure 6C, 6D), which verified that primary cilia were indispensable for Gli2. Western blot detected the expressions of Ac-α-tubulin and Gli2, and further confirmed these results (Figure 6E, 6F).These results suggest that Hedgehog signal and primary cilia are very important for osteoblast differentiation, and Gli2 anchors in primary cilia. High glucose will damage the cilia, blocking of Hedgehog signal and inhibiting osteoblast differentiation. ICA protects primary cilia from high glucose damage, and thus plays a therapeutic effect on impaired osteoblasts differentiation.

**Figure 6.**
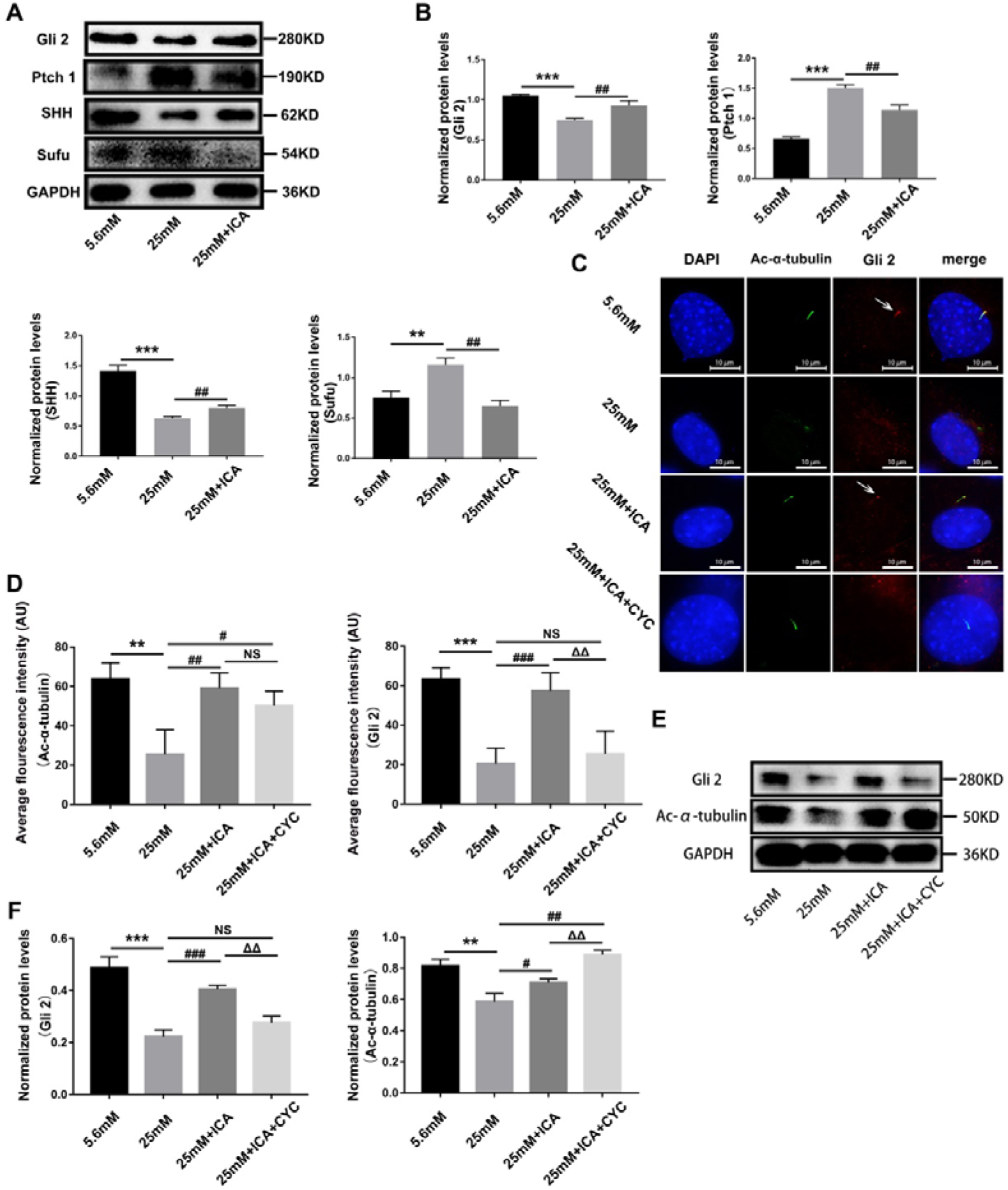
ICA protects primary cilia from high glucose damage and promotes primary cilia localization of Gli2, thus activates the Hedgehog signaling pathway. MC3T3-E1 cells were induced in NC group, HG group and HG+ICA group for 3 days, extracted total cell protein. (A) Western blot detected Hedgehog signaling proteins Gli2, Ptch1, SHH and Sufu, (B) quantitative analysis the protein expression level in (A). MC3T3-E1 cells were induced in NC group, HG group, HG+ICA group and HG+ICA+CYC group for 3 days, (C) immunofluorescence observed primary cilia and Gli2 location, (D) quantitative analysis average fluorescence intensity of Ac-α-tubulin and Gli2 in (C). Extracted total cell protein, (E) Western blot detected Gli2 and Ac-α-tubilin. (F) quantitative analysis the protein expression level in (E). Data were expressed as mean±SEM and were processed by SPSS.22 Student’s t test and one-way ANOVA. *P<0.05, **P<0.01 and ***P<0.001, compared with NC group. ^#^P<0.05, ^##^P<0.01 and ^###^P<0.001, compared with HG group. ^Δ^P<0.05, ^ΔΔ^P<0.01 and ^ΔΔΔ^P<0.001, compared with HG+ICA group.

## Results 7. High glucose inhibits mitochondrial activity, but ICA relieves the inhibition by reducing ROS production

Mitochondrial oxidative phosphorylation produces a large amount of ATP for cell energy supply and is involved in the cell signaling process of energy metabolism. Long-term diabetic hyperglycemia causes the increase of reactive oxygen species (ROS), which can damage mitochondria and thus damage the redox balance of cells^[61]^. Peroxisome proliferator-activated receptor γ coactivator1α (PGC-1α) coordinates the transcription and replication of nuclear and mitochondrial genomes to control mitochondrial biogenesis and maturation. Gain-and loss-of function studies of PGC-1α almost regulate mitochondrial biogenesis, oxidative phosphorylation (OXPHOS), mitochondrial protein levels, and activated nuclear transcription factors that coordinated the upstream input and downstream target genes, such as Nuclear respiratory factor-1 (NRF-1) and mitochondrial transcription factor A (TFAM) through cAMP/CREB, AMPK and other signaling cascades^[62]^. Firstly, we detected the mitochondria-related protein and found the expression of PGC-1α, NRF-1 and TFAM were inhibited significantly in diabetes group SD rats (Fig. 7A, 7B). BMSC, primary OB and MC3T3-E1 cells were induced in NC group, HG group and HG+ICA group, respectively, and Mitotracker was used for immunofluorescence detection of mitochondrial activity after incubation. The results confirmed that, compared with NC group, mitochondrial activities of BMSC, primary OB and MC3T3-E1 cells was significantly reduced in HG group, adding ICA to high glucose medium remarkably increased mitochondrial activity (Fig. 7C, 7D). MC3T3-E1 cells total proteins were extracted, western blot results confirmed the expressions of PGC-1α, NRF-1 and TFAM were inhibited in the HG group, but the addition of ICA significantly rescued the expressions of these three proteins in the high glucose environment (Figure 7E, 7F). To verify whether the protection of ICA on mitochondrial activity is related to the inhibition of ROS production, we induced three kinds of osteoblasts, and then collected cell lysates, DCFH-DA (2’,7’-Dichlorodihydrofluorescein diacetate) probe to detect ROS content. Compared with NC group, high glucose led to a significant increase in ROS in osteoblasts, while ICA could significantly reduce ROS production in high glucose environment. At the same time, we used mitoTempo, a proved mitochondrion-specific scavenger of ROS, as a drug control to inhibit ROS production of osteoblast (Figure 7G). The results showed that the inhibitory effect of ICA on osteoblast ROS production was similar to MitoTempo. The above results suggest that high glucose can damage mitochondrial energy metabolism homeostasis and impair mitochondria activity, while ICA can counteract the damage by inhibiting ROS production.

**Fig 7.**
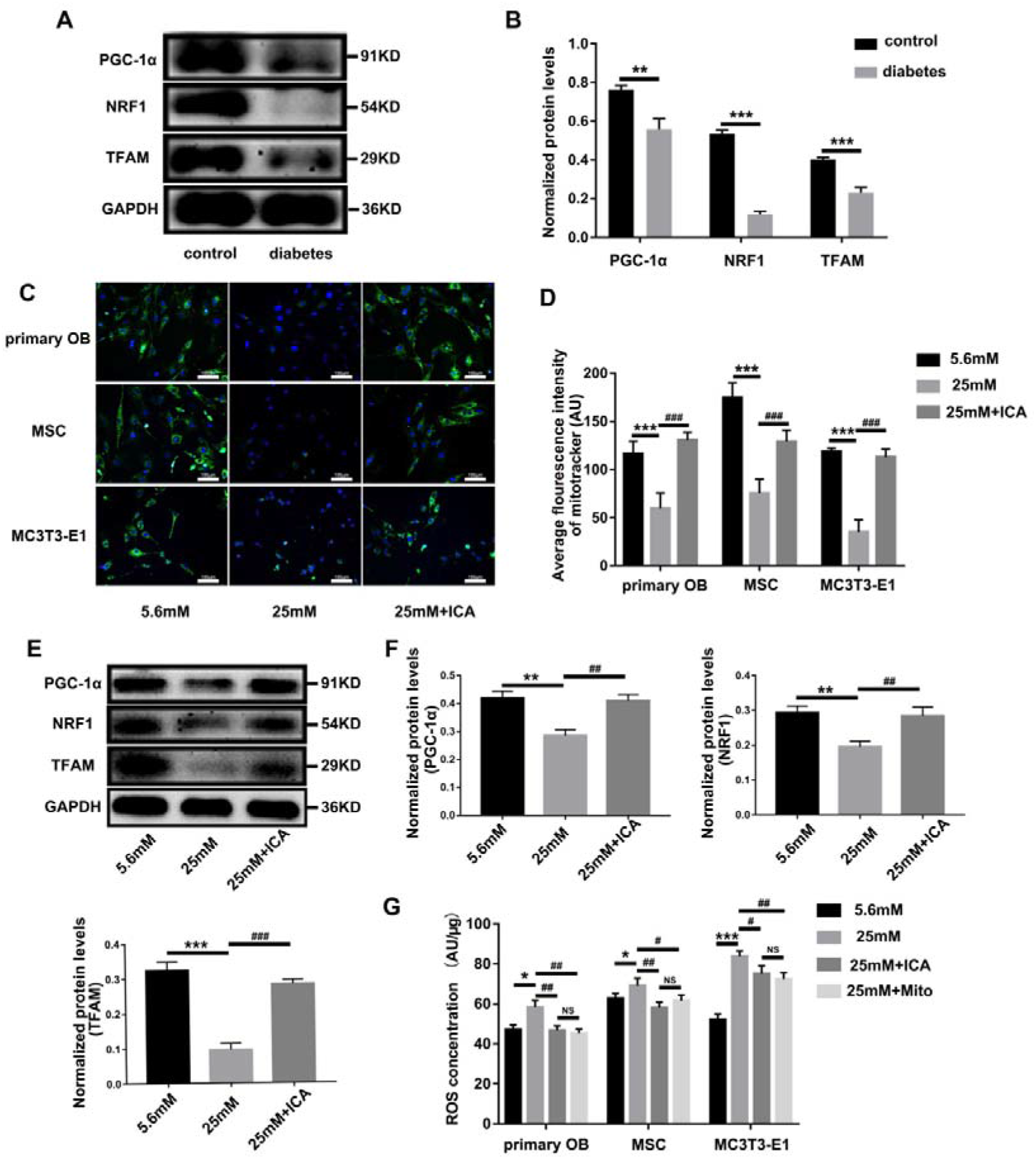
ICA rescues mitochondrial activity from high glucose damage by inhibiting ROS production. (A) Western blot detected the expression of PGC-1α, NRF-1 and TFAM in the femoral protein, (B) Quantitative analysis the level of protein expression in (A). Three kinds of osteoblast cells were induced in NC group, HG group and HG+ICA group for 3 days, (C) immunofluorescence image of Mitotracker staining active mitochondria (green), DAPI staining the nucleus (blue), scale bar: 100μm. (D) Quantitative analysis average fluorescence intensity of (C). (E) Extract total cell protein of MC3T3-E1, Western blot detected PGC-1α, NRF-1 and TFAM, (F) quantitative analysis the protein expression level in (E). (G) The cell lysate of three kinds of osteoblasts was collected, DCFH-DA probe detected ROS content.

**Fig 8.**
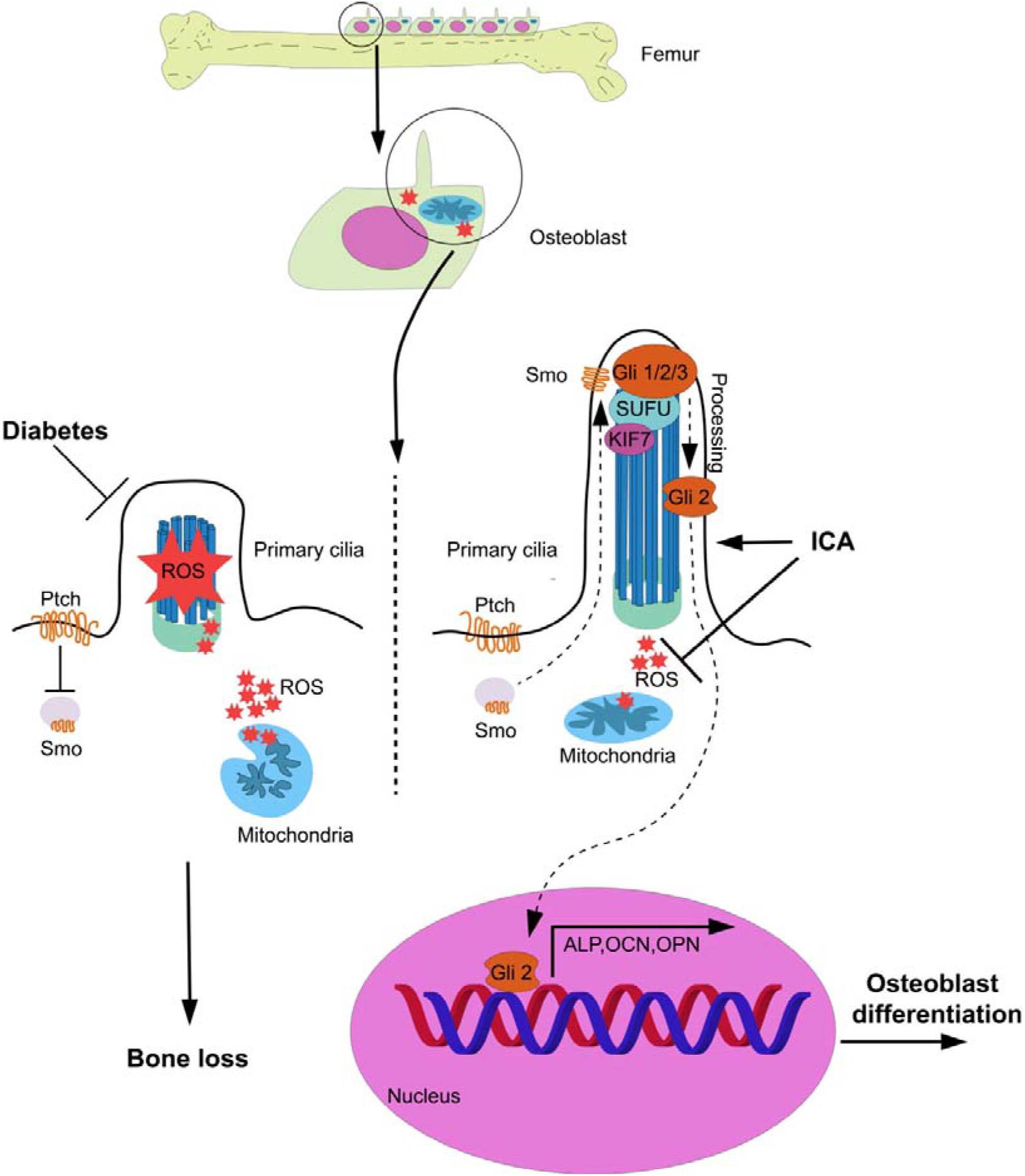
Scheme of the mode of ICA inhibits mitochondrial ROS production to rescue osteoblast differentiation and mineralization by activate cilia/Gli2/Osteocalcin signaling pathway.

## Discussion

Primary cilia are isolated protrusions formed on the surface of mammalian cells, a large number of signal molecules aggregate on the membrane surface or in the cavity to sense the glucose concentration, regulate the secretion of pancreatic islet cells, and mediate calcium ion signals^[34]^. In recent years, more and more studies have found that the occurrence and development of diabetes are closely related to the damage of primary cilia^[63, 64]^, and the primary cilia can be used as an indicator of diabetes risk and treatment targets^[31, 35]^. Our findings confirmed that metabolic disorders contributed by diabetes caused high levels of ROS, which could damage the primary cilia of osteoblasts and inhibit Hedgehog signaling. As a native compound, ICA had antioxidant biological activity and could counteract the damage of ROS to primary cilia, whereby promoted the anchoring of Gli2 on the primary cilia and restored the Hedgehog signal to promote differentiation and mineralization of osteoblasts. The exact mechanism of diabetes induced bone loss is complicated. The reason may be that AGEs, ROS, pro-inflammatory cytokines and adipokines produced by diabetes metabolic disorders form a glycotoxic intracellular environment, which may suppress the proliferation and differentiation of bone cells and destroy bone microstructure and mechanical strength^[6]^. We have a question, does primary cilia dysfunction precede or is it secondary to diabetes? Most of the current researches on primary cilia-related diabetes have focused on primary cilia regulating hormone secretion in pancreatic islet cells. However, we believe that the systemic metabolic disorders caused by progressive diabetes not only damage the primary cilia in pancreas, but also damage the primary cilia in other tissues, directly causing diabetes-related bone, nerve, brain, retina and other complications.

Primary cilia regulate bone development and homeostasis through signal pathways such as BMP-2/Smad4, cAMP/PKA/CREB, Wnt/β-catenin, PI3K/AKT/eNOS^[35, 65–67]^. Mammalian Hedgehog signaling is entirely dependent on the primary cilia, and the two coevolved and adapted to each other to ensure that key signaling molecules are concentrated in the cilia for rapid response to low levels of ligands^[58]^. A large number of studies have confirmed that Hedgehog signaling can couple chondrogenesis with osteogenesis, and regulates bone mass after birth. Some bone diseases (such as skull abnormalities, polydactyly) are considered to be related to abnormal Hedgehog signaling^[68]^. Our previous research on the intraflagellar transport protein IFT80 has found that IFT80 involves in regulating bone development by primary cilia/Hedgehog signal. By targeting to interfere with the synthesis of osteoblast IFT80, we found that the biogenesis of primary cilia were impaired, osteoblast differentiation and mineralization were inhibited. The Smoothened agonists can rescue defects of osteogenesis^[69]^. In this study, we confirmed that the primary cilia in femoral tissues of diabetic SD rats were reduced, and the expression of Hedgehog signaling ligand molecule SHH and nuclear transcription factor Gli2 were significantly decreased as well. This is directly related to the reduction of primary cilia expression caused by high glucose, and impaired primary cilia/Gli2/osteocalcin signal cascades lead to osteoblast differentiation and mineralization block.

Abnormal phenotypes, such as reduced glucose utilization and increased fatty acid consumption, lead to the accumulation of toxic intermediates in diabetic patients ^[70]^. For example, excessive ROS can damage DNA, lipids, endoplasmic reticulum, peroxisomes, mitochondria and other cellular components and organelles^[71]^. Mitochondria not only participate in the synthesis of ATP, but also metabolize fatty acid, lactic acid and ketone body. Mitochondria are the main organelles that generate superfluous ROS^[72]^, which damages mitochondria giving rise to impaired oxidation capacity, increased ROS production, and mitochondrial decoupling^[73, 74]^. Studies on heterotaxy patients and zebrafish mitochondrial-related gene knockout models have confirmed the reduction or dysfunction of mitochondria leads to the elongation of primary cilia, that is, the abundance of mitochondria is negatively correlated with the length of primary cilia, resulting in a ciliopathy phenotype^[75]^, but the impact of excessive ROS on the primary cilia still needs more in-depth research. Our research results confirmed that under high glucose environment, osteoblasts produced a large amount of ROS, which caused to swell and rupture mitochondria, imbalanced oxidative phosphorylation and reduced primary cilia. As a native antioxidant against ROS, ICA rescued the energy metabolism of mitochondria and biogenesis of primary cilia, facilitated osteoblast differentiation and mineralization, suggesting that ICA can be used as a natural drug for the treatment of diabetes induced bone loss.

## Materials and Methods

Study approval: this study has been approved by the Ethics Committee of Affiliated Hospital of Hubei University for Nationalities, and patients and their families are informed of this study and have signed informed consent.

### 4.1. Reagents and antibodies

Icariin (Chengdu Plant Pharmaceutical Factory, purity>98%, Chengdu, China), Cyclopamine (1623, Bristol, UK), α-MEM medium, Fetal Bovine Serum (Gibco), Alizarin Red (Solarbio), Mito-Tracker Green (C1048, Beyotime), alkaline phosphatase kit (NPP substrate-AMP buffer method), Glucose measuring kit (Glucose oxidase method) (BIOSINO BIO-TECHNOLOGY), OCN ELISA kit (MEB1616, Mengbio), DCFH-DA(E004) (Nanjing Jiancheng, China), PGC-1α (1:1000, D162041), TFAM (1:1000, D154208), NRF1(1:1000, D261978) (Sangon Biotech, Shanghai), β-glycerophosphate, dexamethasone, ascorbic acid, acetylated α-tubulin antibody (1:1000, T6793), γ-tubulin antibody (1:1000, T3320), cetylpyridinium chloride (Sigma, USA), Penicillin-Streptomycin Solution (Hyclone), OPN (1:1000, 225952-1-AP, Proteintech), ALP (1:1000, A5111), SHH (1:1000, A5115) (Selleckchem), collagen □ (1:1000, ab255809), OCN (1:1000, ab133612), Runx2 (1:1000, ab54868) (abcam, UK), Ptch1 (1:1000, C53A3), Sufu (1:1000, C54G2), HRP-linked goat anti-rabbit IgG antibody ((1:5000, #7074) and HRP-linked goat anti-mouse IgG antibody (1:5000, #7076) (CST, USA), Gli2 (1:1000, YX3016), GAPDH (1:1000, YT5052, ImmunoWay), Alexa Fluor 568 conjugated anti-rabbit antibody, Alexa Fluor 647 conjugated anti-mouse antibody (Invitrogen, USA).

### 4.2. T1DM SD rats model construction

All studies and operations were performed on rats are according to the procedures approved by Institutional Animal Care and Use Committee of Chongqing Medical University. SD rats aged 8 weeks purchased from experimental Animal Center of Chongqing Medical University. Streptozotocin (STZ, Sangon Biotech, Shanghai) was dissolved into a sodium citrate buffer (50mM, pH=4.5) at a final concentration of 100mg/mL. We first established a type 1 diabetes model by intraperitoneally injecting SD rats at a single large dose STZ (1mL/kg, diabetes group) or sodium citrate buffer (1mL/kg, vehicle control group)^[36]^. After a week, SD rats random blood glucose concentration higher than 250 mg/dL (13.89mmol/L) could be defined as diabetes. Following kept diet SD rats for 90 days to study the effects of diabetes on bone.

### 4.3. Diabetic cell model construction *in vitro*

#### 4.3.1. Primary osteoblasts isolation, culture and osteoblast differentiation induction

Refered to the previous report^[76]^, in a sterile environment, bone marrow mesenchymal stem cells (BMSC) and osteoblasts derived from newborn rat calvarian (primary OB) were isolated from 3-4 days old euthanized rats and were plated in α-MEM medium supplemented with 10% fetal bovine serum (Gibco) and 1% Penicillin-Streptomycin solution (Hyclone) at 37°C, 5% CO_2_ humidified incubator. Selected the cells at passage 3 to seed into 6-well plates at density of 1×10^5^ cells per well. When confluency reached 90%, cells were induced by osteoblast differentiation α-MEM medium (containing 10% FBS, 1% Penicillin-Streptomycin solution, 10mM β-glycerophosphate, 10^−8^M dexamethasone and 50μg/ml ascorbic acid).

#### 4.3.2. Cell model construction

Osteoblasts were induced by osteoblast differentiation α-MEM medium supplemented with D-(+)-glucose to final glucose concentration of 25 mM (HG group) or normal osteoblast differentiation α-MEM medium (NC group), respectively. Moreover, to study the effects of ICA on osteoblast differentiation, ICA was supplemented at final concentrations of 40μmol/L into 25mM group (25mM+ICA). To study the effects of ICA on Hedgehog signaling, Cyclopamine (CYC) was supplemented at final concentrations of 5μmol/L into 25mM+ICA group (25mM+ICA+CYC), and to study the effects of ICA on ROS, Mito-Tempo was supplemented at final concentrations of 5μmol/L into 25mM group (25mM+Mito). After 3 days of induction, total protein was extracted, immunofluorescence was performed, and ROS was determined. After 21 days of induction, mineralized nodules were detected.

#### 4.3.3. Alizarin Red staining mineralized nodules

Osteoblasts were induced for 3 days, removed medium, washed with PBS for 3 times, fixed with 4% cold paraformaldehyde for 10 min, then discarded, washed with PBS for 3 times, stained with 0.2% alizarin red solution (pH 8.3) at room temperature, washed with ddH_2_O for 2 times, and scanned stained cells. Then cells were destained with 10% (W/V) cetylpyridinium chloride for 30 min, the 100μL decolorization solution was transferred to a 96-well plate and optical density (A value) was measured at the wavelength of 562 nm.

#### 4.3.4. Intracellular ROS determination

Osteoblasts were induced for 3 day and collected cell lysates, DCFH-DA (2’,7’-Dichlorodihydrofluorescein diacetate) probe and H_2_O_2_ were diluted at 1:1000. The measurement system consisted of 4μl protein, 46μl ddH_2_O, 100μl diluted DCFH-DA and 100μl diluted H_2_O_2_ solution. Avoiding light and incubating for 30 minutes at 37 ° C, fluorescence intensity was detected according to FITC fluorescence procedures.

#### 4.3.5. ALP activity analysis

Refered to Xu et al. methods^[76]^, ALP activity was determined by ELISA kit (Sigma) according to the manufacturer’s instructions. Briefly, extracted total cell protein after 3 days of osteoblast differentiation induction, the measurement system consisted of 200μl working solution and 4μl cell protein solution. Mixed well, incubated at 37°C for 1 minute and measured the A value at the wavelength of 410 nm every 1 minute, following calculated the rate of change of the A value per minute 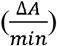. ALP activity is determined according to the formula 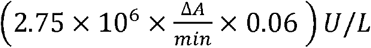.

### 4.4. Immunofluorescence

#### 4.4.1. Primary cilia immunofluorescence

MC3T3-E1 cells were plated on sterile glass coverslips at density of 4 × 10^4^cells/well in 24-well plate. When confluency reached 70%, cells were treated for 3 days. Discarded medium, washed 3 times with pH=7.4 PBS buffer, fixed with 4% paraformaldehyde for 10 minutes, fixed cells were permeabilized with 0.05% Triton X-100 and then washed 3 times with PBS. Incubated with 5% BSA to block non-specific antigen sites for 60 minutes, and then incubated with antibodies of primary cilia structural proteins Ac-α-tubulin antibody and γ-tubulin antibody overnight at 4□ followed by Alexa Fluor568-conjugated anti-rabbit and Alexa Fluor647-conjugated anti-mouse antibodies were used as secondary antibody. Leica DM4000 microscope observed and photographed the cells, ten felds per coverslip (three coverslips per group) were randomly selected.

#### 4.4.2. Mitochondrial Mito-Tracker Green immunofluorescence

Mito-Tracker Green and DAPI were diluted by serum-free medium at 1:5000 and 1:1000, respectively, and incubated at 37 □ in advance. BMSC, primary OB and MC3T3-E1 cells were plated on sterile glass coverslips, respectively, at density of 4 × 10^4^cells/well in 24-well plate. When confluency reached 70%, cells were treated for 3 days. Discarded medium, washed 3 times with serum-free medium, supplement with 500μl preheating Mito-Tracker Green solution and incubated at 37 □ for 30 minutes, discarded medium, washed 3 times, added 200μl DAPI solution and incubated at 37 □ for 40 minutes, Leica DM4000 microscope observed and photographed the cells, ten felds per coverslip (three coverslips per group) were randomly selected.

### 4.5. Immunohistochemistry

Femur and tibia tissues dissected from SD rats of vehicle control group and diabetes group were fixed using 4% cold paraformaldehyde for 48 hours and decalcified in 10% EDTA (pH 7.4) for 60 days at room temperature. After being frozen, the bone tissues were sawed along the largest section of the sagittal plane, longitudinal sections (5 μm thick) were collected on charged glass slides for H&E staining and immunofluorescence. The spleens and pancreases of each group were surgically removed and fixed in 4% paraformaldehyde for 48 hours, then paraffin embedded, section for H&E staining and immunofluorescence.

### 4.6. Micro-computed tomography (Micro-CT)

Micro-CT was performed to evaluate bone mechanical properties and microarchitecture, 4% paraformaldehyde fixed left femur was scanned using a Micro-CT scanner (SCANCO Medical AG, vivaCT40). Trabecular bone thickness parameters are calculated by distance transformation (Direct-No Model) and surface triangulation (TRI-Plate Model) methods. BV, TV, BV/TV, Tb.Th, Tb.N, Tb.Sp, BS, BS/BV, SMI were calculated from the region of interest (ROI). X-ray tube: micro focus tube supporting conical X-ray beam, focal diameter of 5μm. The date was analysed by instrument software.

### 4.7. Western blot

Osteoblasts were induced for 3 days, collected cell total protein and measured the protein concentration with BCA kit (Beyotime, China). The protein was denatured in 6× loading buffer and separated in 10% SDS-PAGE gel. Transferred the protein to the PVDF membrane in a buffer solution of 0.192 mol/L glycine, 0.25 mol/L Tris and 20% methanol. After blocking with 5% defatted milk-TBST for 1 hours, the membrane was incubated with primary antibody overnight at 4°C, and then incubated with horseradish peroxidase (HRP)-conjugated goat anti-rabbit/mouse IgG antibody for 1 hours. Detected target protein signal with Western Bright ECL chemiluminescence reagent, and visualized was performed using BIO RAD ChemiDoc™ Touch Imaging System (USA). GAPDH was used as a loading control, the gray value of protein band were measured using Image J software, The experiment was performed in triplicate.

### 4.8. Transmission Electron Microscopy (TEM)

Blocked SD rat pancreas and spleen approximately 1 mm^3^ in size were double-fixed with 2.5% glutaraldehyde and 1% osmium acid at 4□ overnight. Dehydration was done in grades of ethanol (50%, 70%, 90%) and 90% acetone, and specimens were embedded in Epon812 epoxy resin, ultrathin sections (50 nm) were mounted on copper grids and contrasted with uranyl acetate and lead acetate, transmission electron microscope(JEM-1400PLUS, JEOL, Japan) observed and photoed.

### 4.9. Statistical Analysis

All data are presented as mean ± SEM. (N ≥ 3). Student’s t-test are used to comparison between two groups and one-way ANOVA, followed by Tukey’s multiple comparison test are analyzed by grouped samples. P<0.05 was used as a threshold for statistical significance. GraphPad Prism was used for these analyses.

## Acknowledgments

Research reported in this publication was supported by research grants from Chongqing Yuzhong district the first plan of scientific research in 2020 (Grant No. 20200112), Chongqing Graduate Science and Technology innovation Projects in 2019 (Grant No. CYS19204), Doctoral Program of Higher Education of China (Grant No. 20125503120015), the Natural Science Foundation of Chongqing in China (Grant No.cstc2014jcyjA10024) and the Education Commission of Chongqing in China (Grant No.CY170402). The content is solely the responsibility of the authors and does not necessarily represent the official views of the financially supported government.

## Author contributions

Chang dong W. and Jie L. designed the research and analyzed the data. Jie L. Xiaoyan D, Xiangmei W, Maorong W, Shengyong Y, Jie X, Qian C, Mengxue L, Xianjun L, Changdong W. designed, performed experiments and analyzed the data.

## Reference

[1] Q&A: Key points for IDF Diabetes Atlas 2017. Diabetes research and clinical practice, 2018, 135(235-236.

[2] KDIGO 2020 Clinical Practice Guideline for Diabetes Management in Chronic Kidney Disease. Kidney international, 2020, 98(4s): S1–s115.

[3] Klein K R, Buse J B. The trials and tribulations of determining HbA(1c) targets for diabetes mellitus. Nature reviews Endocrinology, 2020, 16(12): 717–730.

[4] Johnell O, Kanis J A. An estimate of the worldwide prevalence and disability associated with osteoporotic fractures. Osteoporosis international : a journal established as result of cooperation between the European Foundation for Osteoporosis and the National Osteoporosis Foundation of the USA, 2006, 17(12): 1726–1733.

[5] Shahen V A, Gerbaix M, Koeppenkastrop S, et al. Multifactorial effects of hyperglycaemia, hyperinsulinemia and inflammation on bone remodelling in type 2 diabetes mellitus. Cytokine & growth factor reviews, 2020, 55(109-118.

[6] Napoli N, Chandran M, Pierroz D D, et al. Mechanisms of diabetes mellitus-induced bone fragility. Nature reviews Endocrinology, 2017, 13(4): 208–219.

[7] Jiang Y, Zhang Y, Chen W, et al. Correction to: Achyranthes bidentata extract exerts osteoprotective effects on steroid-induced osteonecrosis of the femoral head in rats by regulating RANKL/RANK/OPG signaling. Journal of translational medicine, 2021, 19(1): 208.

[8] Ma H, Wang X, Zhang W, et al. Melatonin Suppresses Ferroptosis Induced by High Glucose via Activation of the Nrf2/HO-1 Signaling Pathway in Type 2 Diabetic Osteoporosis. Oxidative medicine and cellular longevity, 2020, 2020(9067610.

[9] Savelli B, Li Q, Webber M, et al. RedoxiBase: A database for ROS homeostasis regulated proteins. Redox Biol, 2019, 26(101247.

[10] Ahmad W, Ijaz B, Shabbiri K, et al. Oxidative toxicity in diabetes and Alzheimer’s disease: mechanisms behind ROS/RNS generation. Journal of biomedical science, 2017, 24(1): 76.

[11] Dugan L L, You Y H, Ali S S, et al. AMPK dysregulation promotes diabetes-related reduction of superoxide and mitochondrial function. The Journal of clinical investigation, 2013, 123(11): 4888–4899.

[12] Vestergaard P. Discrepancies in bone mineral density and fracture risk in patients with type 1 and type 2 diabetes—a meta-analysis. Osteoporosis International, 2007, 18(4): 427–444.

[13] Kurra S, Fink D A, Siris E S. Osteoporosis-associated fracture and diabetes. Endocrinology and metabolism clinics of North America, 2014, 43(1): 233–243.

[14] Khosla S, Bilezikian J P, Dempster D W, et al. Benefits and risks of bisphosphonate therapy for osteoporosis. The Journal of clinical endocrinology and metabolism, 2012, 97(7): 2272–2282.

[15] Kearns A E, Khosla S, Kostenuik P J. Receptor activator of nuclear factor kappaB ligand and osteoprotegerin regulation of bone remodeling in health and disease. Endocrine reviews, 2008, 29(2): 155–192.

[16] Neer R M, Arnaud C D, Zanchetta J R, et al. Effect of parathyroid hormone (1-34) on fractures and bone mineral density in postmenopausal women with osteoporosis. The New England journal of medicine, 2001, 344(19): 1434–1441.

[17] Cosman F, Crittenden D B, Adachi J D, et al. Romosozumab Treatment in Postmenopausal Women with Osteoporosis. The New England journal of medicine, 2016, 375(16): 1532–1543.

[18] Khosla S, Hofbauer L C. Osteoporosis treatment: recent developments and ongoing challenges. The lancet Diabetes & endocrinology, 2017, 5(11): 898–907.

[19] Khosla S, Burr D, Cauley J, et al. Bisphosphonate-associated osteonecrosis of the jaw: report of a task force of the American Society for Bone and Mineral Research. Journal of bone and mineral research : the official journal of the American Society for Bone and Mineral Research, 2007, 22(10): 1479–1491.

[20] Shane E, Burr D, Abrahamsen B, et al. Atypical subtrochanteric and diaphyseal femoral fractures: second report of a task force of the American Society for Bone and Mineral Research. Journal of bone and mineral research : the official journal of the American Society for Bone and Mineral Research, 2014, 29(1): 1–23.

[21] Wang L, Li Y, Guo Y, et al. Herba Epimedii: An Ancient Chinese Herbal Medicine in the Prevention and Treatment of Osteoporosis. Current pharmaceutical design, 2016, 22(3): 328–349.

[22] Wisanwattana W, Wongkrajang K, Cao D Y, et al. Inhibition of Phosphodiesterase 5 Promotes the Aromatase-Mediated Estrogen Biosynthesis in Osteoblastic Cells by Activation of cGMP/PKG/SHP2 Pathway. Front Endocrinol (Lausanne), 2021, 12(636784.

[23] Simpson E R, Clyne C, Rubin G, et al. Aromatase--a brief overview. Annual review of physiology, 2002, 64(93-127.

[24] Wu Y, Cao L, Xia L, et al. Evaluation of Osteogenesis and Angiogenesis of Icariin in Local Controlled Release and Systemic Delivery for Calvarial Defect in Ovariectomized Rats. Sci Rep, 2017, 7(1): 5077.

[25] Zhou L, Poon C C, Wong K Y, et al. Icariin ameliorates estrogen-deficiency induced bone loss by enhancing IGF-I signaling via its crosstalk with non-genomic ERα signaling. Phytomedicine : international journal of phytotherapy and phytopharmacology, 2021, 82(153413.

[26] Indran I R, Liang R L, Min T E, et al. Preclinical studies and clinical evaluation of compounds from the genus Epimedium for osteoporosis and bone health. Pharmacology & therapeutics, 2016, 162(188-205.

[27] Kim B, Lee K Y, Park B. Icariin abrogates osteoclast formation through the regulation of the RANKL-mediated TRAF6/NF-κB/ERK signaling pathway in Raw264.7 cells. Phytomedicine : international journal of phytotherapy and phytopharmacology, 2018, 51(181-190.

[28] Yang A, Yu C, Lu Q, et al. Mechanism of Action of Icariin in Bone Marrow Mesenchymal Stem Cells. Stem cells international, 2019, 2019(5747298.

[29] Yuan X, Serra R A, Yang S. Function and regulation of primary cilia and intraflagellar transport proteins in the skeleton. Annals of the New York Academy of Sciences, 2015, 1335(1): 78–99.

[30] Delaine-Smith R M, Sittichokechaiwut A, Reilly G C. Primary cilia respond to fluid shear stress and mediate flow-induced calcium deposition in osteoblasts. FASEB journal : official publication of the Federation of American Societies for Experimental Biology, 2014, 28(1): 430–439.

[31] Kluth O, Stadion M, Gottmann P, et al. Decreased Expression of Cilia Genes in Pancreatic Islets as a Risk Factor for Type 2 Diabetes in Mice and Humans. Cell Rep, 2019, 26(11): 3027–3036.e3023.

[32] Lee S Y, Long F. Notch signaling suppresses glucose metabolism in mesenchymal progenitors to restrict osteoblast differentiation. The Journal of clinical investigation, 2018, 128(12): 5573–5586.

[33] Ehnert S, Sreekumar V, Aspera-Werz R H, et al. TGF-beta1 impairs mechanosensation of human osteoblasts via HDAC6-mediated shortening and distortion of primary cilia. J Mol Med (Berl), 2017, 95(6): 653–663.

[34] Hughes J W, Cho J H, Conway H E, et al. Primary cilia control glucose homeostasis via islet paracrine interactions. Proc Natl Acad Sci U S A, 2020, 117(16): 8912–8923.

[35] Shi W, Gao Y, Wang Y, et al. The flavonol glycoside icariin promotes bone formation in growing rats by activating the cAMP signaling pathway in primary cilia of osteoblasts. J Biol Chem, 2017, 292(51): 20883–20896.

[36] Cheng R X, Feng Y, Liu D, et al. The role of Na(v)1.7 and methylglyoxal-mediated activation of TRPA1 in itch and hypoalgesia in a murine model of type 1 diabetes. Theranostics, 2019, 9(15): 4287–4307.

[37] Furman B L. Streptozotocin-Induced Diabetic Models in Mice and Rats. Current protocols in pharmacology, 2015, 70(5.47.41-45.47.20.

[38] Eizirik D L, Björklund A, Cagliero E. Genotoxic agents increase expression of growth arrest and DNA damage--inducible genes gadd 153 and gadd 45 in rat pancreatic islets. Diabetes, 1993, 42(5): 738–745.

[39] Goodpaster B H, Sparks L M. Metabolic Flexibility in Health and Disease. Cell metabolism, 2017, 25(5): 1027–1036.

[40] Schmidt A M. Highlighting Diabetes Mellitus: The Epidemic Continues. Arteriosclerosis, thrombosis, and vascular biology, 2018, 38(1): e1–e8.

[41] Cole J B, Florez J C. Genetics of diabetes mellitus and diabetes complications. Nature reviews Nephrology, 2020, 16(7): 377–390.

[42] Morgan E F, Unnikrisnan G U, Hussein A I. Bone Mechanical Properties in Healthy and Diseased States. Annual review of biomedical engineering, 2018, 20(119-143.

[43] Verdelis K, Lukashova L, Atti E, et al. MicroCT morphometry analysis of mouse cancellous bone: intra-and inter-system reproducibility. Bone, 2011, 49(3): 580–587.

[44] Nishiyama K K, Campbell G M, Klinck R J, et al. Reproducibility of bone micro-architecture measurements in rodents by in vivo micro-computed tomography is maximized with three-dimensional image registration. Bone, 2010, 46(1): 155–161.

[45] Bala Y, Zebaze R, Seeman E. Role of cortical bone in bone fragility. Current opinion in rheumatology, 2015, 27(4): 406–413.

[46] Cooper D M, Kawalilak C E, Harrison K, et al. Cortical Bone Porosity: What Is It, Why Is It Important, and How Can We Detect It? Curr Osteoporos Rep, 2016, 14(5): 187–198.

[47] Harada S, Rodan G A. Control of osteoblast function and regulation of bone mass. Nature, 2003, 423(6937): 349–355.

[48] Wei J, Karsenty G. An overview of the metabolic functions of osteocalcin. Reviews in endocrine & metabolic disorders, 2015, 16(2): 93–98.

[49] Komori T. Functions of Osteocalcin in Bone, Pancreas, Testis, and Muscle. International journal of molecular sciences, 2020, 21(20):

[50] Nijhuis A W, Van Den Beucken J J, Jansen J A, et al. In vitro response to alkaline phosphatase coatings immobilized onto titanium implants using electrospray deposition or polydopamine-assisted deposition. Journal of biomedical materials research Part A, 2014, 102(4): 1102–1109.

[51] Sengottuvelan A, Balasubramanian P, Will J, et al. Bioactivation of titanium dioxide scaffolds by ALP-functionalization. Bioactive materials, 2017, 2(2): 108–115.

[52] Halling Linder C, Englund U H, Narisawa S, et al. Isozyme profile and tissue-origin of alkaline phosphatases in mouse serum. Bone, 2013, 53(2): 399–408.

[53] García-Hernández A, Arzate H, Gil-Chavarría I, et al. High glucose concentrations alter the biomineralization process in human osteoblastic cells. Bone, 2012, 50(1): 276–288.

[54] Ibsen C J, Birkedal H. Modification of bone-like apatite nanoparticle size and growth kinetics by alizarin red S. Nanoscale, 2010, 2(11): 2478–2486.

[55] Anvarian Z, Mykytyn K, Mukhopadhyay S, et al. Cellular signalling by primary cilia in development, organ function and disease. Nature reviews Nephrology, 2019, 15(4): 199–219.

[56] Moore E R, Zhu Y X, Ryu H S, et al. Periosteal progenitors contribute to load-induced bone formation in adult mice and require primary cilia to sense mechanical stimulation. Stem cell research & therapy, 2018, 9(1): 190.

[57] Kluth O, Stadion M, Gottmann P, et al. Decreased Expression of Cilia Genes in Pancreatic Islets as a Risk Factor for Type 2 Diabetes in Mice and Humans. Cell Rep, 2019, 26(11): 3027–3036 e3023.

[58] Bangs F, Anderson K V. Primary Cilia and Mammalian Hedgehog Signaling. Cold Spring Harbor perspectives in biology, 2017, 9(5):

[59] Nygaard M B, Almstrup K, Lindbæk L, et al. Cell context-specific expression of primary cilia in the human testis and ciliary coordination of Hedgehog signalling in mouse Leydig cells. Sci Rep, 2015, 5(10364.

[60] Santos N, Reiter J F. A central region of Gli2 regulates its localization to the primary cilium and transcriptional activity. Journal of cell science, 2014, 127(Pt 7): 1500–1510.

[61] Ruegsegger G N, Creo A L, Cortes T M, et al. Altered mitochondrial function in insulin-deficient and insulin-resistant states. The Journal of clinical investigation, 2018, 128(9): 3671–3681.

[62] Dorn G W, 2nd, Vega R B, Kelly D P. Mitochondrial biogenesis and dynamics in the developing and diseased heart. Genes Dev, 2015, 29(19): 1981–1991.

[63] Gerdes J M, Christou-Savina S, Xiong Y, et al. Ciliary dysfunction impairs beta-cell insulin secretion and promotes development of type 2 diabetes in rodents. Nat Commun, 2014, 5(5308.

[64] Volta F, Scerbo M J, Seelig A, et al. Glucose homeostasis is regulated by pancreatic β-cell cilia via endosomal EphA-processing. Nat Commun, 2019, 10(1): 5686.

[65] Liang W, Lin M, Li X, et al. Icariin promotes bone formation via the BMP-2/Smad4 signal transduction pathway in the hFOB 1.19 human osteoblastic cell line. Int J Mol Med, 2012, 30(4): 889–895.

[66] Zhai Y K, Guo X Y, Ge B F, et al. Icariin stimulates the osteogenic differentiation of rat bone marrow stromal cells via activating the PI3K-AKT-eNOS-NO-cGMP-PKG. Bone, 2014, 66(189-198.

[67] Baron R, Kneissel M. WNT signaling in bone homeostasis and disease: from human mutations to treatments. Nat Med, 2013, 19(2): 179–192.

[68] Ohba S. Hedgehog Signaling in Skeletal Development: Roles of Indian Hedgehog and the Mode of Its Action. International journal of molecular sciences, 2020, 21(18):

[69] Yang S, Wang C. The intraflagellar transport protein IFT80 is required for cilia formation and osteogenesis. Bone, 2012, 51(3): 407–417.

[70] Zorzano A, Liesa M, Palacín M. Role of mitochondrial dynamics proteins in the pathophysiology of obesity and type 2 diabetes. The international journal of biochemistry & cell biology, 2009, 41(10): 1846–1854.

[71] Ji Y, Chae S, Lee H K, et al. Peroxiredoxin5 Controls Vertebrate Ciliogenesis by Modulating Mitochondrial Reactive Oxygen Species. Antioxidants & redox signaling, 2019, 30(14): 1731–1745.

[72] Tsushima K, Bugger H, Wende A R, et al. Mitochondrial Reactive Oxygen Species in Lipotoxic Hearts Induce Post-Translational Modifications of AKAP121, DRP1, and OPA1 That Promote Mitochondrial Fission. Circulation research, 2018, 122(1): 58–73.

[73] Boudina S, Sena S, Theobald H, et al. Mitochondrial energetics in the heart in obesity-related diabetes: direct evidence for increased uncoupled respiration and activation of uncoupling proteins. Diabetes, 2007, 56(10): 2457–2466.

[74] Boudina S, Bugger H, Sena S, et al. Contribution of impaired myocardial insulin signaling to mitochondrial dysfunction and oxidative stress in the heart. Circulation, 2009, 119(9): 1272–1283.

[75] Burkhalter M D, Sridhar A, Sampaio P, et al. Imbalanced mitochondrial function provokes heterotaxy via aberrant ciliogenesis. The Journal of clinical investigation, 2019, 129(7): 2841–2855.

[76] Xu J, Deng X, Wu X, et al. Primary cilia regulate gastric cancer-induced bone loss via cilia/Wnt/β-catenin signaling pathway. Aging, 2021, 13(6): 8989–9010.

